# Biological signatures of history: Examination of composite biomes and Y chromosome analysis from da Vinci-associated cultural artifacts

**DOI:** 10.64898/2026.01.06.697880

**Authors:** Harinder Singh, Seesandra V. Rajagopala, Rebecca Hart, Pille Hallast, Mark Loftus, Rosana Wiscovitch-Russo, Cody R. K. Conrad, David S. Thaler, Guadalupe Piñar, Karina C. Åberg, Rossella Lorenzi, José A Lorente, Jesse H. Ausubel, Thomas P. Sakmar, Rhonda K. Roby, Charles Lee, Norberto Gonzalez-Juarbe

## Abstract

Cultural heritage objects can accumulate DNA from materials, environments, and repeated human contact, but biomolecular profiling of such items is constrained by nondestructive sampling requirements, ultra-low biomass, and high contamination risk. Here we present a minimally invasive workflow that integrates gentle swab collection, low- input whole-metagenome sequencing, taxonomic profiling, and Y-chromosome analyses to recover “biological signatures of history” from Renaissance-era artwork and archival correspondence associated with ancestors of Leonardo da Vinci. Across artifacts, we recovered heterogeneous mixtures of microbial and eukaryotic DNA (including bacteria, fungi, plants, and viruses) consistent with composite “biomes” that reflect differences in substrate, storage, conservation treatments, and handling. Multivariate comparisons show reproducible sample-to-sample separations. In parallel, we assessed human Y-chromosome signal using a panel of ∼90,000 phylogenetically informative markers and partial Y-STR profiling where feasible. Across multiple independent swabs from Leonardo da Vinci-associated items, the obtained Y chromosome marker data suggested assignments withing the broader E1b1/E1b1b clade. However, the control samples also indicate mixed contributions consistent with modern handling and other sources. Together, these data demonstrate the feasibility as well as limitations of combining metagenomics and human DNA marker analysis for cultural heritage science, providing a baseline workflow for future conservation science studies and hypothesis-driven investigations of provenance, authentication and handling history.

## Introduction

Cultural heritage objects (including drawings, manuscripts, and archival correspondence) can accumulate biological residues over time, including microbial DNA, environmental DNA (deposited during storage and transport), and human DNA (introduced through repeated handling)^1^. Because these cultural objects are often unique and fragile, sampling must be minimally invasive. Consequently, recovered DNA is typically ultra-low biomass and highly susceptible to modern contamination^2,3^. These constraints make it challenging to distinguish signals plausibly associated with an object’s long-term history from those introduced by recent handling, storage environments, and laboratory processing.

Advances in next generation sequencing (NGS) and taxonomic profiling have enabled broad surveys of DNA recovered from heritage materials, with practical applications in conservation science, including monitoring biodeterioration risk and characterizing microbial communities associated with storage and treatment conditions^4–7^. For surface sampling in particular, interpretation must remain conservative. Indeed, DNA profiles are composites by default (substrate, conservation materials, dust / aerosols, and multiple human contacts) and low signal-to-noise means analytical choices and control design can strongly influence which taxa appear credible. As a result, claims about provenance, geolocation, or historical events require authentication, replication, and contamination aware controls.

In parallel, there is interest in whether human DNA recovered from cultural objects can provide information about the creators, owners, or handlers. Unfortunately, human DNA on swabbed surfaces is commonly fragmented, present at very low abundance (relative to microbial DNA), and often represents mixtures from multiple contributors, limiting genealogical inference^8^. The Y chromosome is a plausible exploratory target because it is inherited along the paternal line and structured into well characterized haplogroups. Recent advancement in human Y chromosome assemblies and variant catalogs have further refined haplogroup frameworks^9,10^. Nevertheless, robust Y haplogroup assignment still depends on sufficient, internally consistent marker coverage and careful analyses of DNA mixtures and contamination.

Here, we present an exploratory, minimally invasive workflow that combines a gentle double swab sampling approach^11^, low-input whole-metagenome sequencing without host DNA removal, and taxonomic profiling. We coupled this with a feasibility analysis of male specific human DNA using Y chromosome markers and partial Y-STR profiling where possible. We apply this framework to a red chalk drawing attributed to Leonardo da Vinci (1452-1519), the “Holy Child” (dated to ∼1472-1476)^12^ alongside archival correspondence from Frosino di Ser Giovanni da Vinci (an ancestor of Leonardo da Vinci) to Luca del Sera from the Archivio di Stato di Prato^8^ and comparator drawings by other master artists (Filippino Lippi, Andrea Sacchi, and Charles J. Flipart) (**Fig. 1A**).

**Figure 1:**
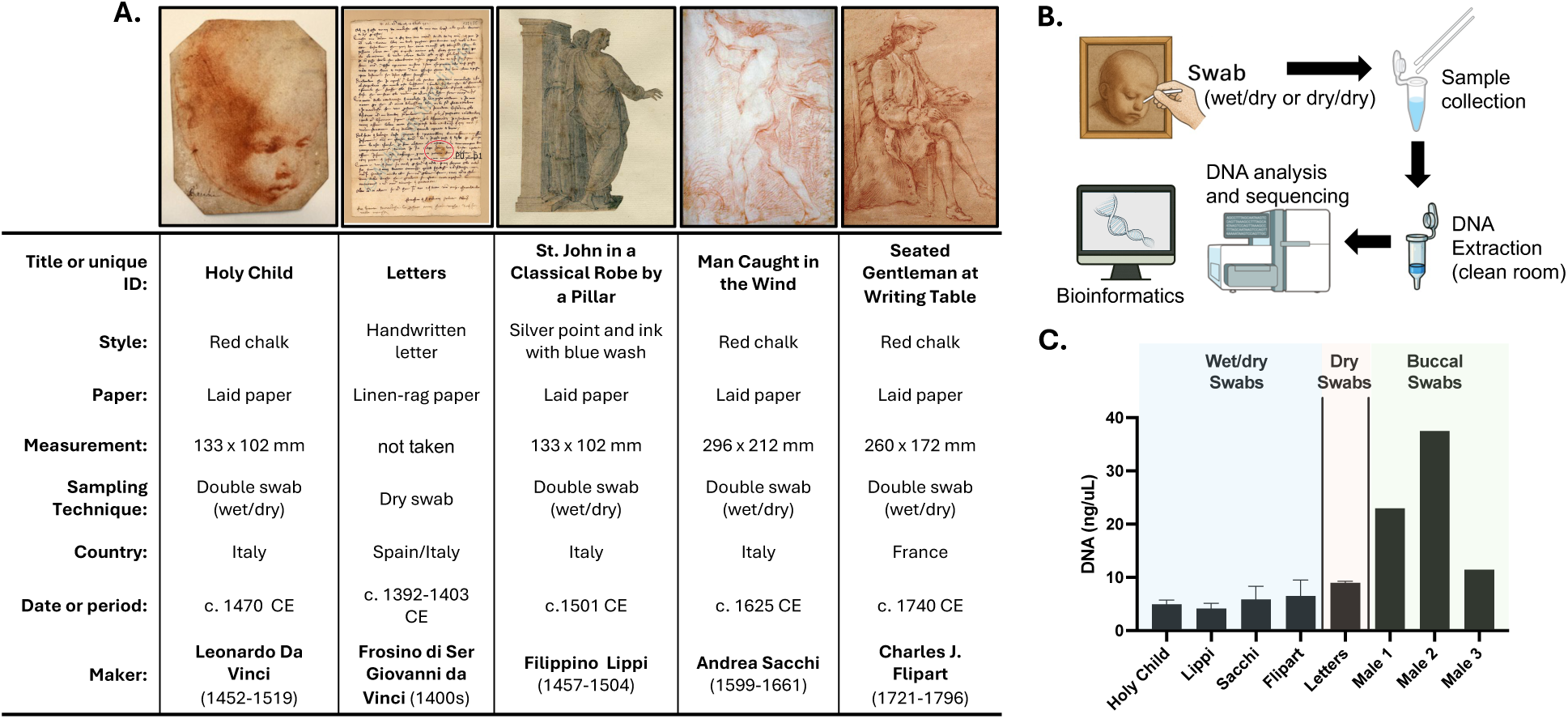
Isolation and amplification of ancient DNA from art obtained by the double-swab technique. **a)** Detailed description and pictures of the artwork or correspondence from Leonardo da Vinci, Frosino di Ser Giovanni da Vinci, Filippino Lippi, Andrea Sacchi and Charles J. Flipart. **b)** Illustration of platform used for sample acquisition, DNA isolation, next generation sequencing and bioinformatics. **c)** Quantification of DNA isolated ng/µL from each cultural artifact of male controls with regards to the method of collection used.

Our aims are deliberately constrained by the properties of low biomass surface DNA: (a) to describe the composite, multi domain biological material detectable on these cultural objects under a standardized minimally invasive workflow, (b) to test whether samples show reproducible differences in recovered community composition under the same pipeline, and (c) to assess the feasibility and limitations of extracting interpretable Y chromosome signal from metagenomic short reads using contemporary controls to contextualize potential modern contributions. Taken together, we present a minimally-invasive sampling and analysis framework intended to establish a baseline characterization of the biological material detectable on selected Leonardo-associated cultural artifacts, while explicitly delimiting which historical or provenance interpretations are supported by the data.

## Results

### DNA recovery from cultural heritage artifacts and comparator samples

DNA recovered from cultural heritage surfaces is typically low in quantity and may be fragmented and/or chemically damaged^3^. In addition, surface swabs are inherently susceptible to modern contamination and mixed contributions from handlers and environments. Previous research has demonstrated that DNA can be recovered using a range of approaches (ranging from minimally-invasive dry swabbing to more invasive wet vacuuming approaches), with higher yields generally associated with more destructive techniques^11^.

Using previously described double swab collection procedures (wet/dry or dry/dry, depending on object constraints)^11^, we sampled artworks and archival correspondence (**Fig. 1B**). DNA was extracted from each swab and total DNA was quantified (**Fig. 1C**). Of note, the dry/dry swab technique provided comparable DNA yield as that of the wet/dry. Given the low-input nature of these extracts, we prepared whole-metagenome sequencing libraries directly from recovered DNA without attempting any host (human) DNA removal. Then, we performed library enrichment as part of the library preparation protocol prior to sequencing.

### Multi domain DNA is detected across cultural artifacts

Sequencing-based surveys of cultural artifacts can recover heterogeneous biological profiles shaped by multiple factors including object substrate, conservation materials, storage environments, and handling history^13^. Consistent with this expectation, read mapping-based analyses assigned DNA to an abundance of diverse organisms (**Fig. 2A**). Using a stringent classification threshold, the most abundant assignments fell into broad groups including plants, viruses, animals, bacteria and fungi (**Fig. 2B**). We then applied a secondary scaffold/contig based classification using a more relaxed threshold to capture additional low abundance signal. This approach recovered the same broad composition (**Fig. S1**), but increased the number of taxa reported and therefore required more cautious interpretation at the individual taxon level.

**Figure 2:**
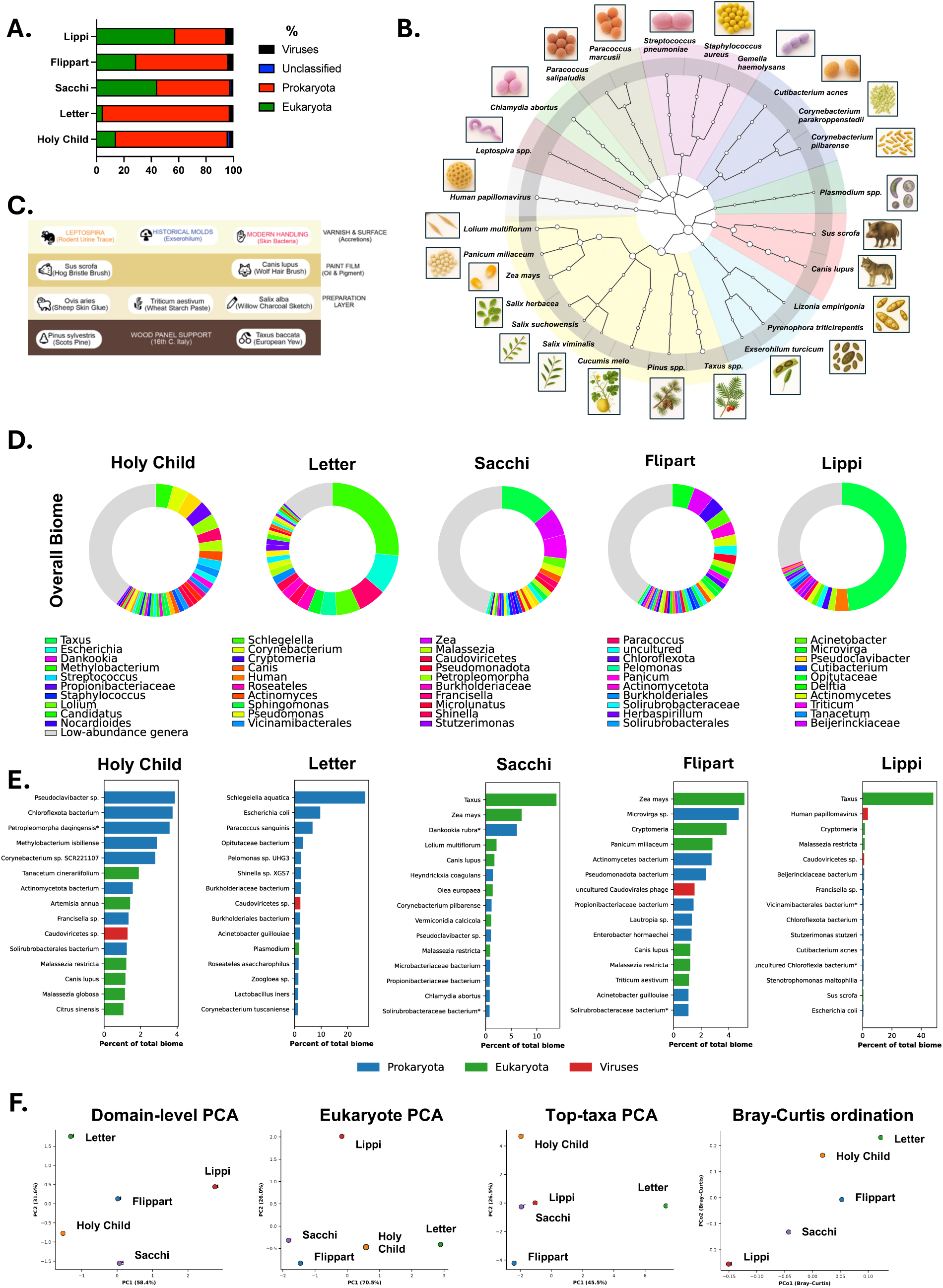
Distinct biome profiles are present in ancient cultural artifacts. **A)** Percent abundance of viruses, prokaryotes, eukaryotes and unclassified organisms in Holy Child, Letter, Sacchi, Lippi and Flipart. **B)** Taxonomic rank classification for highly abundant curated organisms. **C)** Metagenomic stratigraphy. **D)** Total biome genus classification with top 50 genera with a unique color-identifier. **E)** Bar plots for the top 15 most abundant taxa with no forensic correction for each artifact. **F)** Multivariate comparisons of total biome composition across datasets using principal component analysis (PCA) and distance-based ordination.

Within the plant associated assignments, taxa detected across samples included *Lolium multiflorum*, *Panicum miliaceum*, *Zea mays*, *Salix* spp., *Cucumis melo*, *Pinus* spp. and *Taxus* spp. (**Fig. 2B**; **Fig. S2**). Scaffold level classification also included *Citrus* spp. (**Fig. S2**) with *Citrus sinensis* showing its highest relative abundance in the “Holy Child” sample (**Fig. S2**). Because plant DNA on artworks can originate from numerous sources (*e.g*., ambient dust/pollen, paper and plant derived materials, storage environments, conservation treatments and handling), these detections are interpreted primarily as description of recovered sequence assignments rather than as definitive evidence of a specific historical origin.

Animal associated assignments were dominated by *Sus scrofa* and *Canis lupus* (**Fig. 2B**), with the highest relative abundance observed (in descending order) in Sacchi, Flipart, the “Holy Child” and Lippi, with little to no signal in the archival letter (**Fig. 2B**). *Sus scrofa* was also detected in Lippi, Sacchi and the “Holy Child” in the scaffold-based analysis (**Fig. S2**). As with plants, animal DNA on cultural objects may derive from more recent environmental deposition or modern handling and should not be conclusively assigned to a specific source without additional validation.

Fungal assignments under the stringent cutoff were dominated by taxa including *Lizonia emperigonia*, *Pyrenophora tritici-repentis* and *Exserohilum turcicum* (**Fig. 2B**), with the strongest signal in the Flipart sample. Under the relaxed scaffold classification, additional fungal taxa were detected, including *Vermiconidia*, *Malassezia* and *Stereum* (**Fig. S2**). Of note, *Malassezia* spp. are common human skin associated fungi and their presence on cultural objects is consistent with modern handling rather than artifact-specific historical signal^14^.

One result requiring particular caution was the detection of sequences assigned to *Plasmodium* spp. in the archival letter. This appeared at substantially higher relative abundance (10 times) than in other artifacts (**Fig. 2B; Fig. S2**). Because low biomass metagenomic datasets are vulnerable to spurious taxonomic assignments and contamination, this finding is treated as preliminary until supported by targeted validation (*e.g*., read level verification across multiple *Plasmodium* loci).

Viral assignments were dominated by sequences classified as human papillomavirus and bacteriophages (**Fig. 2B; Fig. S1**). Across domains, several detected taxa have been reported previously in studies of artwork and cultural artifacts as surface contaminants, tools used to draw or paint, or organisms associated with human contact (**Fig. 2C**). We interpret these results as composite biological signatures detected by our workflow. Attribution to provenance, geography, or specific historical exposures would require additional authentication.

### Bacterial profiles dominate classified reads

Bacterial assignments constituted the largest fraction of classified reads across samples (**Fig. 2A**). Several high abundance taxa were consistent with modern sources including skin associated organisms such as *Cutibacterium acnes* (**Fig. 2B**). Additional bacterial assignments (e.g., *Leptospira* spp.; **Fig. 2B**) were also observed. When analyses were restricted beyond the most ubiquitous skin associated taxa, the relative bacteriall composition differed across artifacts (**Fig. S1**). At the broader level, the combined multi-domain composition (prokaryotes, eukaryotes, and viruses together) also differed in relative proportions across objects (**Fig. 2D-E**).

To test whether these sample-to-sample differences were detectable in multivariate space, we compared total biome composition across datasets using principal component analysis (PCA) and distance-based ordination. Relative abundance tables were generated from classifications mapped to the NCBI Taxonomy lineage and summarized at multiple levels: broad domains, selected eukaryotic subgroups, and sets of high abundance taxa. Features were Z-score standardized prior to PCA, and Bray–Curtis dissimilarities were used for ecological ordination. Across these analyses, samples separated in ordination space (**Fig. 2F**), suggesting differences in recovered community composition. Further validation with expanded negative controls, replication, and sensitivity analyses would be required to distinguish artifact associated signal from technical and modern contamination.

### Detectable Human DNA enables feasibility focused Y-chromosome analysis

Across datasets, reads mapping to the human genome varied by sample (**Fig. S3**), consistent with differences in the amount of human material recovered by swabbing. To evaluate feasibility of recovering male specific human genetic signal from swab derived metagenomic data, and to assess whether any signal could reflect contributors involved in the creation, ownership, or subsequent handling of the objects, we focused on reads mapping to the male-specific region of the Y chromosome. The Y chromosome is inherited along the paternal line and contains phylogenetically informative markers defining Y haplogroups with characteristic geographic distributions^9,10^.

We used the Illumina short read sequence data from 16 samples, including three independent swabs from the “Holy Child”, swabs from ten letters written by Leonardo’s ancestor Frosino, three commercially acquired drawings of similar style to the “Holy Child”, and comparator art pieces from Flipart, Sacchi and Lippi. To contextualize modern contributions, we included multiple controls. These included swabs from commercially available art (from the same historical period), four human controls (3 males and 1 female), swabs from the frame in which the “Holy Child” was stored, and environmental swabs from the sample collection rooms in the USA and Italy (**Fig. 3A**). Of note, DNA extraction from all samples was performed solely by female researchers.

**Figure 3:**
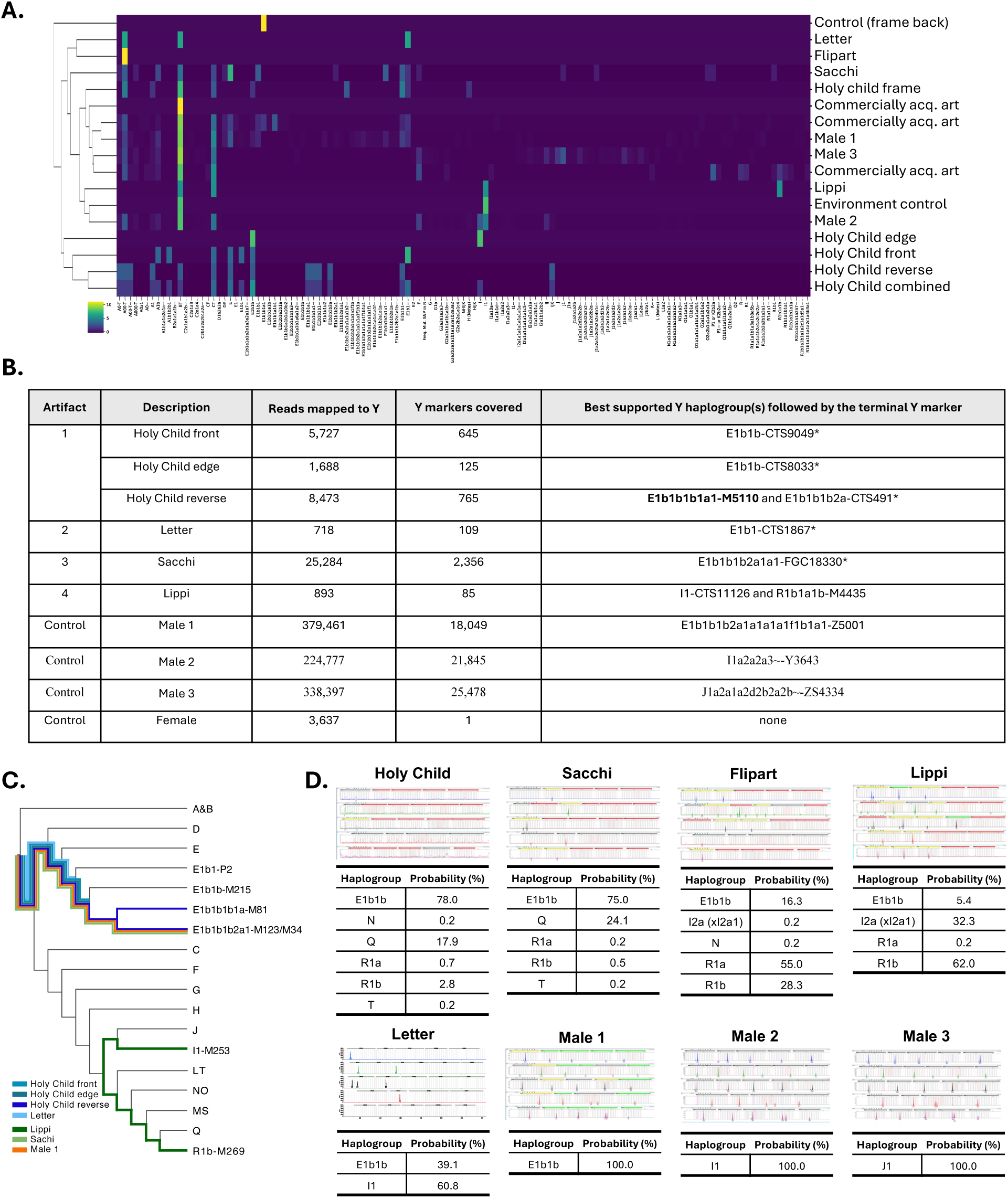
Y-chromosome haplogroup inference from short read sequences and Y STR profiling supports haplogroup identification within ancient cultural artifacts associated with Leonardo da Vinci. **A)** Hierarchically clustered heat map of haplogroup markers using derived ISOGG marker counts. Colors indicate z-score magnitude (high - yellow, low - dark blue). **B)** Top supported Y haplogroups for samples with sufficient data. (* - haplogroup indistinguishable, at the current resolution, from that inferred for the control men and female). **C)** The supported phylogenetic placement of sequenced artefacts and the male researcher**. D)** Y-STR (Y-filer) based % probability for haplogroup.

For haplogroup inference, we queried a panel of ∼90,000 phylogenetically informative Y-chromosome markers. For each sample, we determined the allelic states of markers present in the read data and inferred the best supported haplogroup(s) (**Fig. 3A-B**). Y marker coverage was sparse overall (0–2,356 markers per sample; mean 261), and only 5 of the 16 samples had sufficient informative coverage for a more stable haplogroup call. As expected, the female scientist served as a negative control for Y-chromosome signal (**Fig. 3B**).

Across samples labeled as Leonardo da Vinci-associated (three swabs from the “Holy Child” and the aggregate letter dataset), the best supported haplogroup assignments fell within the broader E1b1 clade (**Fig. 3B-C**). The front and edge swabs from the “Holy Child” supported E1b1b sublineages (E1b1b-CTS9049 and E1b1b-CTS8033), the aggregate letters supported an upstream E1b1 branch (E1b1-CTS1867). The reverse swab from the “Holy Child” supported two closely related E1b1b1b sublineages (E1b1b1b1a1-M5055 and E1b1b1b2a-CTS491) (**Fig. 3B**). Control Male 1 also carried a E1b1b1b haplogroup (E1b1b1b2a1a1a1a1f1b1a1-Z5001), although with additional phylogenetic divergence at the available resolution. Control Male 2 and Male 3 showed haplogroups I1a2a2a3-Y3643 and J1a2a1a2d2b2a2b-ZS4334, respectively (**Fig. 3B**).

We also observed isolated reads supporting ancestral state markers in the “Holy Child” and single marker support for haplogroup “I” and macrohaplogroup IJK (*i.e*., I. J1, R1). Given sparse coverage, these single read observations cannot be resolved confidently and may reflect noise, mis-mapping, or mixture. Overall, the repeated recovery of E1b1/E1b1b class assignments across multiple Leonardo-associated samples is compatible with a shared Y-chromosomal signal (**Fig. 3B, C**).

Among the remaining objects, Sacchi showed best support for a Y lineage E1b1b1b2a1a1-FGC18330 (**Fig. 3B, C**). Lippi showed limited support for I1 and R1b1a1b lineages, both common in Europe, though the number of informative markers precluded confident placement into a more specific subclade.

### Partial Y STR profiles support feasibility of male marker recovery

To further assess feasibility of human male marker recovery, we generated Y-STR profiles (27 loci; Yfiler Plus) for selected samples (including the “Holy Child”, the Letters, Sacchi, Flipart, Lippi, and males controls 1-3). Complete 27 locus profiles were obtained only for the three male controls. In contrast, the “Holy Child” yielded partial calls at 6 loci, and three letters yielded partial calls at 7 loci. Other artifacts produced partial and variable call sets (*i.e*., Sacchi 2 loci; Flipart 9 loci; Lippi 16 loci), and several loci showed more than one allele, consistent with mixed male contributions (**Fig. S4**).

Because these artifact derived profiles are partial and often composite, we estimated haplogroup posterior probabilities from Y-STR values using a Bayesian allele frequency approach (**Fig. 3D**)^15^. Under this framework, the “Holy Child” had its highest posterior probability assigned to E1b1b (78.0%), followed by Q (17.9%) and R1b (2.8%). The letters showed split posterior probability between E1b1b (39.1%) and I1 (60.8%). Sacchi showed mixed posterior probability across E1b1b (75.0%), Q (24.1%), and low probability assignments to R1a (0.2%), R1b (0.5%) and T (0.2%). Flipart also showed mixed posterior probabilities across R1a (55.0 %), R1b (28.3%) and smaller probabilities for E1b1b (16.3%), I2a (xI2a1) (0.2%), N (0.2%) lineages. Lippi showed higher posterior probabilities for R1b (62.0%) and I2a (xI2a1) (32.3%) than E1b1b (5.4%) or R1a (0.2%). Male controls 1-3 showed 100% posterior probability to E1b1b, I1 and J1, respectively. These Y STR results are broadly consistent with the short read haplogroup assignments, while also reinforcing key limitations of surface derived material from ancient historical artifacts.

## Discussion

This study evaluates what can (and cannot) be learned from minimally invasive swab sampling of cultural heritage objects combined with low input whole metagenome sequencing and downstream taxonomic and Y chromosome analyses. Across Leonardo associated drawings and archival correspondence, comparator artworks and contemporary controls, we recovered heterogeneous mixtures of non-human DNA (microbial and eukaryotic) and, in a subset of samples, sparse male-specific human DNA signals. Together, these results support the feasibility of detecting multi domain biological material on cultural objects under a standardized low input workflow while underscoring the central constraint of surface derived DNA. Recovered profiles are composites by default and highly susceptible to modern contributions and analytical artifacts.

### Composite biome profiles

Microbiome based profiling has been proposed as a tool for conservation monitoring and, more speculatively, for provenance or geolocation inference^13^. In our dataset, taxonomic classification identified diverse bacteria, fungi, plants, animals, and viruses across artifacts. The profiles represent composite biological signatures shaped by a combination of object substrate, conservation materials, storage environments and human handling. Because many taxa plausibly originate from recent contact and environmental deposition, individual organism detection should not be treated as direct evidence of origin, artistic intent, or a specific historical setting without additional authentication and contamination aware controls.

Several plant and environmental taxa detected in this study are broadly compatible with sources expected for works on paper in European contexts (e.g., dust/pollen deposition, paper- and plant derived materials, and storge related inputs). Certain non-human eukaryote DNA may help us to understand the artifact composition, possible materials used, the environment and geology of the pieces obtained during the Renaissance in Florence and other areas of Europe (ca. 1400–1550 CE). For example plants such as *Lolium multiflorum* (commonly known as Italian ryegrass, *Panicum miliaceum* - one of the oldest domesticated crops in human history)^16^, were present in Italy in the 1400s and 1500s. Riparian species such as Salix spp. were abundant along the Arno River and were commonly used for basketry, bindings, scaffolding, and charcoal production within artisanal workshops^17^. Similarly, coniferous taxa such as Pinus spp. were widely exploited in Tuscany for panel supports, framing, and construction, while pine resins and pitch were routinely incorporated into sealants and varnishes used in artistic production^18^. The unique presence of Citrus spp. in the “Holy Child” may provide a direct link to historical context. The Medici family were known as patrons of art, gardens, and experimental horticulture in Tuscany^19^. Medici villas housed collections of exotic plants, with citrus trees occupying a central role^20,21^. For the Medici’s, citrus plants were seen as a symbol of dynastic wealth, global connectivity, and scientific curiosity^22^. Interestingly, Leonardo da Vinci is documented by early biographer Giorgio Vasari, as having worked under the patronage of Lorenzo de’ Medici in Florence and spending time in the Medici garden at San Marco, a space devoted to sculpture, artistic training, and humanist exchange^23^.

Environmental taxa detected in datasets may also reflect historical importance. For example, *Alternaria alternata*, is a ubiquitous saprophytic fungus found in soil, dust, and plants. It is also one of the most frequently encountered fungi on paper, wooden objects, and painted surfaces, and is widely recognized as a contributor to biodeterioration^24^. Similarly, the identification of *Leptospira* species is consistent with the ecology of Renaissance Florence, where rodent infestations, livestock circulation, and damp storage areas would have facilitated environmental deposition of soil- and water-associated spirochetes^25^. Similarly, the detection of *Chlamydia abortus*, an ovine and caprine pathogen, is likewise plausible in a regional context marked by extensive sheep husbandry for wool production^26^. Finally, *Plasmodium* spp. represents an organism of clear historical relevance: malaria was endemic across multiple regions of Italy during the Renaissance, including lowland and marsh-adjacent areas of Tuscany^27^. Furthermore, recent analyses of Medici family burials have confirmed malarial infection in several individuals^28^. Collectively, these taxa illustrate the mixture of environmental and regionally plausible organisms that can accumulate on cultural artifacts during their inception time or during extended storage. Future studies should combine the current approach with other Omic analyses such as proteomics, metabolomics and transcriptomics, to further define, with higher resolution, the composite biomes of the samples^1,29–31^.

### Y-chromosome signal feasibility

We also tested whether short read metagenomic data from swabbed objects can support exploratory inference of paternal lineage markers. Y chromosome marker coverage was sparse for most samples, and only a minority of datasets supported more stable haplogroup assignment. This limitation is expected for low template, surface derived human DNA and constrains the strength of lineage inferences.

Across Leonardo associated samples, the best supported assignments often fell withing the broader E1b1/E1b1b clade. This clade is common across the Mediterranean, in parts of central and southern Italy, including Tuscany^32,33^, and therefore its presence, by itself, cannot conclusively distinguish between historical contributors and modern handlers. Importantly, at least one male control also carried an E1b1b (E-M215) haplogroup, and at the resolution enabled by sparse marker coverage, overlap between artifact samples and controls is consistent with mixed contributions.

The Holy Child reverse swab is assigned to the E1b1b1b-M81 branch, which today reaches its highest frequencies in North Africa^34,35^. Outside North Africa, E-M81 occurs at low but detectable frequencies in southern Europe, including Italy and especially Sicily and parts of southern mainland regions, where it is interpreted as reflecting historical North African gene flow across the central Mediterranean. In contrast, a male control associated with the study, carries a downstream lineage within the E-M123/E-M34 branch (subclade Z5001), a near eastern-derived component of E1b1b that shows its highest frequencies in Levantine and neighboring populations and occurs at low frequencies across North Africa and Europe, including Italy^36^. The recovery of an E1b1b1b-M81 lineage from the Holy Child reverse swab, together with basal E1b1b lineages in the “Holy Child” front and edge swabs and in Frosino’s letter, is consistent with the presence of a coherent E1b1b signal on the Leonardo-related artefacts, with more recent handling-derived DNA likely contributing additional Y-chromosomal profiles. Of importance, the separate evaluation of the Y haplogroups in the same samples by Y-STR profiling provides further support for the E1b1b hypothesis as several of the other artifacts had very low percent haplogroup probabilities for E1b1b, reducing the notion that the signal may be solely modern contamination. Thus, the consistent detection of E1b1 lineages across multiple independent swabs, together with a related family document, and the cross validation by Y-STR profiling supports the notion of a shared Y-chromosomal signal linking Leonardo-associated artifacts. However, future studies testing additional Leonardo-associated artifacts such as notebooks, sketches, and paintings, or confirmed living descendants are required to properly conclude that this is the da Vinci lineage^37,38^.

In summary, minimally invasive swab-based sampling, coupled with low input next-generation DNA sequencing, can recover detectable, multi-domain biological material from cultural heritage objects and can support comparative profiling across artifacts and controls. The principal limitation of this study is that low biomass swab metagenomics is extremely sensitive to contamination and to analytical decisions. To enable stronger claims, especially relating to provenance, geolocation, or historical characteristics, future work is needed to help distinguish artifact-associated signal from modern handling. Nevertheless, the current dataset provides a practical baseline for feasibility and conservation-oriented monitoring hypotheses.

## Methods

### Artwork used

The “Holy Child” was provided by art historian Fred R. Kline. The piece of art is a renaissance-era drawing (*c.* 1472-76) on laid paper that measures 5.25 × 4.25 inches, with a blank verso and a watermark of a partial cut-off sun or star symbol, potentially made by Leonardo da Vinci (*c.* 1452-1519). The medium is red chalk and white chalk on uncoated laid paper with corners later cropped; traces of sepia ink framing around the drawing (notable around the right side of drawing); and sepia ink notations and sepia ink spotting at the lower left. The other selected artworks (also from the Fred R. Kline collection) were: “St. John in a Classical Robe by a Pillar”, a silverpoint drawing with ink and blue wash on laid paper by Filippino Lippi; “Man Caught in the Wind”, a red chalk drawing on laid paper by Andrea Sacchi; and the “Seated Gentleman at Writing Table”, a red chalk with white highlights on laid paper by Charles J. Flipart. Letters from Frosino di Ser Giovanni da Vinci, an ancestor of Leonardo da Vinci who lived in Barcelona, Spain, were obtained from the Fondo Datini at the Archivio di Stato di Prato in Prato, Italy.

### Sample collection

Biological samples were collected as previously described in Roby *et al*. (2025)^11^ and others^39,40^. Artifacts from the Fondo Datini at the Archivio di Stato di Prato were sampled using a double-dry swab method, while all other works were sampled using the wet-dry double-swabbing method^11^. The wet swab was used to very gently rub the surface of the area to be sampled while employing a swirling motion. Subsequently, the same surface area was sampled with a second dry swab using the same technique. After allowing the swabs to air dry under conditions that prevent environmental contamination, both swabs were stored in a sterile tube. The dry swab technique was done as the wet-dry technique, without any buffer. Of note, both techniques yielded comparable DNA yields (**Fig. 1C**). Buccal swabs were collected by placing a sterile swab in the cheek are within the mouth and swirling for 15-30 seconds. All living human subjects (3 males and 1 female) were fully informed about the use of genetic information for this study and the related ethical matters. A consent form was signed for the use of the collected material, analysis was done in blind fashion, with all identifiers removed.

### Isolating DNA

DNA from the samples was extracted manually using the Macherey-Nagel NucleoMag-DNA Forensic kit inside a PCR clean room. The kit utilized magnetic beads to bind the DNA. Samples that had two swabs present were processed by removing the swab heads from the swab sticks and placing half of each the wet and the dry swabs together into a tube for DNA extraction. At the elution step, the two factions were then combined to have a single sample. Samples that had a single swab present, were processed by removing the swab head and placing the entire swab into a tube for DNA extraction. After the extraction, the total amount of DNA present was quantified using the Thermo Fisher Nanodrop and Qubit. The ancient cultural artifact samples were quantified using 2µL of sample while the Kline Collection samples were quantified using 1µL of sample.

### Metagenomics

The Illumina sequencing libraries were prepared using the Twist Bioscience library preparation Kit, which enzymatically fragments genomic DNA to a target size range, performs end-repair, dA-tailing, adapter ligation, and library amplification for 10 cycles, using unique dual index (UDI) primers. Library quality was assessed using the Agilent Bioanalyzer with the DNA high-sensitivity chip, and quantification was performed via qPCR using the KAPA library quantification kit and a QuantStudio 12K instrument. Sequencing was performed on the Illumina NovaSeq X Plus platform, generating 2×150 bp paired-end reads with an average depth of approximately 50 million reads per sample.

### Assembly and Pre-processing

Metagenomic sequencing reads were assembled using metaSPAdes v3.10^41^, utilizing 32 threads and a memory limit of 250 GB to ensure robust graph traversal. Following assembly, scaffold headers were standardized to include sample-specific prefixes (e.g., “all ”) using SeqKit^42^. A Simplified Annotation Format (SAF) file was subsequently generated to facilitate downstream feature quantification.

### Read Mapping and Adaptive Filtering

To estimate scaffold coverage and abundance, the original paired-end reads were mapped back to the final assembly using Bowtie2^43^. An index was constructed using bowtie2-build, and alignment was performed in “very-sensitive” mode with end-to-end alignment parameters (--very-sensitive, --end-to-end) to maximize sensitivity. To minimize spurious alignments, strict pairing criteria were enforced by prohibiting mixed (--no-mixed) and discordant (--no-discordant) alignments. The --no-unal flag was employed to exclude unaligned reads from the output.

Post-alignment processing was conducted using SAMtools^44^. Resulting SAM files were converted to BAM format, sorted by name for mate-fixing (fixmate -m), and subsequently coordinate-sorted. PCR duplicates were identified and removed using markdup -r. Alignments were strictly filtered to retain only properly paired, primary alignments with a minimum mapping quality (MAPQ) of 30.

### Adaptive Contig Selection

An adaptive selection strategy was applied to the filtered BAM files to retain contigs meeting robust detection thresholds: 1. Primary Thresholds: Minimum breadth of coverage of 40% and a minimum mean depth of 5X. 2. Adaptive Fallback: If no contigs met primary thresholds, criteria were adjusted to the 95th percentile of mean depth and 75th percentile of breadth observed in the sample. 3. Minimum Selection: If the fallback yielded zero contigs, the top 50 contigs by mean depth were selected.

### Feature Quantification

Read abundance was quantified using featureCounts^45^ in paired-end mode (-p). Counting mode was dynamically determined: strict fragment requirements (-p -B -C) were enabled if properly paired reads were detected; otherwise, single-end counting was utilized. All features were quantified with a minimum mapping quality filter of 30 (-Q 30) and un-stranded settings (-s 0). Quantification was performed at two resolutions: ORFs: Coding sequences predicted by Prodigal^46^ were quantified using the GTF format option. Scaffolds: Full contigs were quantified using the SAF format option.

### Taxonomic Assignment

Taxonomic classification of each ORF and contig was performed using the CAT (Contig Annotation Tool) package^47^ with default parameters. The resulting abundance tables were processed using a custom Python 3 workflow utilizing the Pandas library. Taxonomic identifiers were normalized by truncating strings at the first occurrence of colon-like delimiters. Data were aggregated by species identity, summing abundance counts across all sample columns. To reduce noise, the dataset was filtered to exclude species with a cumulative abundance count of less than 10.0 across all samples.

### Y-chromosome haplogroup inference from Illumina sequence data

Reads were mapped using bwa (v0.7.19-r1273)^48^ mem to the T2T-CHM13 (autosomes+X) plus GRCh38 chromosome Y reference. Samtools (v1.21)^49^ was used to obtain mapped read counts, to add mate tags, sort reads by name, and mark duplicates. Bcftools (v1.22 using htslib 1.22.1)^48,49^ mpileup/call was used to obtain allelic read counts across the ∼10.4 Mb of “callable with Illumina” Y regions^50^ for both the invariant and variant sites, using minimum base quality 20 and minimum mapping quality 20. All calls within 5 base pairs of an indel call were removed (bcftools filter -g 5), followed by the removal of indels. All sites were compared against 90,002 phylogenetically informative Y-chromosome markers from ISOGG (www.isogg.org, v15.73), after excluding duplicate coordinates and indels. Both reference and variant alleles (relative to GRCh38 Y) were assessed for phylogenetic informativeness and for the concordance with the topology of the human Y phylogeny. Variant sites from the artefacts were checked for signs of cytosine deamination to uracil (observed as C>T and complementary G>A substitutions), considered an ancient DNA damage signature, and terminal markers consistent with such signature were excluded. Samples were hierarchically clustered using average linkage and Euclidean distance on row-wise z-scored values from derived ISOGG marker counts using seaborn clustermap python package^51^.

### Y-STR analysis

Based on the Quantifiler Trio quantification results, there was extremely low amounts of Y-chromosome in the samples. Due to the low quantity, the samples were concentrated down using an Eppendorf Vacufuge and then rehydrated in 10µl of elution buffer. The Y-chromosome was analyzed using the Thermo Fisher Yfiler Plus kit. The entire 10µl of sample was loaded into the Yfiler Plus reaction and amplified on the Thermo Fisher SimpliAmp Thermal Cycler. After amplification, the samples were then loaded onto the Thermo Fisher 3500xl Genetic Analyzer and the allele calling was done using the GeneMapper software.

## Acknowledgements

We thank Fred Kline and Angela Zimm for the contributions of artwork for this project. We thank the Fondo Datini at the Archivio di Stato di Prato in Prato, Italy for the contributions of cultural artifact samples for this project and in particular Leonardo Meoni, Virginia Barni and Chiara Marcheschi. We thank Drs. Peter Vallone and Erica Romsos of the Applied Genetics Group at the National Institute of Standards and Technology for their essential technical support in conducting the quantitative polymerase chain reaction (qPCR) testing and capillary electrophoresis short tandem repeat (CE STR) typing. We also acknowledge Thomas Huber, Manija Kazmi, Marguerite Mangin and members of the Leonardo Da Vinci DNA Project for support and discussions. We also thank Drs. J. Craig Venter, Karen Nelson and Manolito Torralba for their support and discussions. CL is a scientific advisor for Nabsys, LLC.

## References

1 White, A. E. et al. Ancient DNA and biomarkers from artefacts: insights into technology and cultural practices in Neolithic Europe. Proc Biol Sci 292, 20250092 (2025). 10.1098/rspb.2025.0092

2 Flocco, C. G., Methner, A., Burkart, F., Geppert, A. & Overmann, J. Touching the (almost) untouchable: a minimally invasive workflow for microbiological and biomolecular analyses of cultural heritage objects. Front Microbiol 14, 1197837 (2023). 10.3389/fmicb.2023.1197837

3 Orlando, L. et al. Ancient DNA analysis. Nature Reviews Methods Primers 1, 14 (2021). 10.1038/s43586-020-00011-0

4 Maljkovic Berry, I., et al. Next Generation Sequencing and Bioinformatics Methodologies for Infectious Disease Research and Public Health: Approaches, Applications, and Considerations for Development of Laboratory Capacity. J Infect Dis 221, S292–S307 (2020). 10.1093/infdis/jiz286

5 Marciniak, S. & Perry, G. H. Harnessing ancient genomes to study the history of human adaptation. Nat Rev Genet 18, 659–674 (2017). 10.1038/nrg.2017.65

6 Vilanova, C. & Porcar, M. Art-omics: multi-omics meet archaeology and art conservation. Microb Biotechnol 13, 435–441 (2020). 10.1111/1751-7915.13480

7 Pinar, G. et al. The Microbiome of Leonardo da Vinci’s Drawings: A Bio-Archive of Their History. Front Microbiol 11, 593401 (2020). 10.3389/fmicb.2020.593401

8 Gorden, E. M. et al. Next generation sequencing of STR artifacts produced from historical bone samples. Forensic Sci Int Genet 49, 102397 (2020). 10.1016/j.fsigen.2020.102397

9 Hallast, P. et al. Assembly of 43 human Y chromosomes reveals extensive complexity and variation. Nature 621, 355–364 (2023). 10.1038/s41586-023-06425-6

10 Rhie, A. et al. The complete sequence of a human Y chromosome. Nature 621, 344–354 (2023). 10.1038/s41586-023-06457-y

11 Roby, R. K. et al. Sampling techniques and genomic analysis of biological material from artworks. J Forensic Sci 70, 476–489 (2025). 10.1111/1556-4029.15701

12 Kline, F. R. Leonardo’s Holy Child. (Pegasus Books, 2016).

13 Piñar, G., Poyntner, C., Tafer, H. & Sterflinger, K. A time travel story: metagenomic analyses decipher the unknown geographical shift and the storage history of possibly smuggled antique marble statues. Annals of Microbiology 69, 1001–1021 (2019). 10.1007/s13213-019-1446-3

14 Rahimlou, S., Amend, A. S. & James, T. Y. Malassezia in environmental studies is derived from human inputs. mBio 16, e0114225 (2025). 10.1128/mbio.01142-25

15 Athey, T. W. Haplogroup Prediction from Y-STR Values Using a Bayesian-Allele-Frequency Approach. Journal of Genetic Genealogy, 34–39 (2006).

16 Lu, H. et al. Earliest domestication of common millet (Panicum miliaceum) in East Asia extended to 10,000 years ago. Proc Natl Acad Sci U S A 106, 7367–7372 (2009). 10.1073/pnas.0900158106

17 Gennai, M. et al. The floodplain woods of Tuscany: towards a phytosociological synthesis. Plant Sociology 58 (2021). 10.3897/pls2021581/01

18 The Structural Conservation of Panel Paintings: Proceedings of a Symposium at the J. Paul Getty Museum, 24–28 April 1995. (The Getty Conservation Institute, 1998).

19. Gihring, T. Meet the Medicis: The mad, marvelous family behind the Italian Renaissance, <https://new.artsmia.org/stories/meet-the-medicis-the-mad-marvelous-family-behind-the-italian-renaissance> (2022).

20 Sophia Cabrera, M. J., Carson Sanders, Gabriel McCown & Shauna O’Callaghan. Exploring the Medici Garden of Castello: How Art Holds Power, <https://www.auf-florence.org/exploring-the-medici-garden-of-castello-how-art-holds-power/> (

21 Medici Villas and Gardens in Tuscany, <https://villegiardinimedicei.it/en/medici-villas-and-gardens-of-tuscany/> (2006).

22 (Leonardo da Vinci Art School).

23 Vasari, G. The Life Of Leonardo Da Vinci. (44, 2019).

24 Koul, B. & Upadhyay, H. in Fungi and their Role in Sustainable Development: Current Perspectives (eds Praveen Gehlot & Joginder Singh) 597–615 (Springer Singapore, 2018).

25 Carmichael, A. G. Plague and the poor in Renaissance Florence. (Cambridge University Press, 2014).

26 Santori, D. et al. Biomolecular survey on the main infectious causes of abortion in sheep in the Italian regions of Latium and Tuscany. Open Vet J 14, 1447–1452 (2024). 10.5455/OVJ.2024.v14.i6.12

27 Boualam, M. A. et al. The millennial dynamics of malaria in the mediterranean basin: documenting Plasmodium spp. on the medieval island of Corsica. Front Med (Lausanne) 10, 1265964 (2023). 10.3389/fmed.2023.1265964

28 Fornaciari, G., Giuffra, V., Ferroglio, E., Gino, S. & Bianucci, R. Plasmodium falciparum immunodetection in bone remains of members of the Renaissance Medici family (Florence, Italy, sixteenth century). Trans R Soc Trop Med Hyg 104, 583–587 (2010). 10.1016/j.trstmh.2010.06.007

29 Creydt, M. & Fischer, M. Artefact Profiling: Panomics Approaches for Understanding the Materiality of Written Artefacts. Molecules 28 (2023). 10.3390/molecules28124872

30 Poinar, H. N. et al. Metagenomics to paleogenomics: large-scale sequencing of mammoth DNA. Science 311, 392–394 (2006). 10.1126/science.1123360

31 Marmol-Sanchez, E. et al. Ancient RNA expression profiles from the extinct woolly mammoth. Cell (2025). 10.1016/j.cell.2025.10.025

32 Boattini, A. et al. Uniparental markers in Italy reveal a sex-biased genetic structure and different historical strata. PLoS One 8, e65441 (2013). 10.1371/journal.pone.0065441

33 Capelli, C. et al. Y chromosome genetic variation in the Italian peninsula is clinal and supports an admixture model for the Mesolithic-Neolithic encounter. Mol Phylogenet Evol 44, 228–239 (2007). 10.1016/j.ympev.2006.11.030

34 Cruciani, F. et al. Phylogeographic analysis of haplogroup E3b (E-M215) y chromosomes reveals multiple migratory events within and out of Africa. Am J Hum Genet 74, 1014–1022 (2004). 10.1086/386294

35 Sole-Morata, N. et al. Whole Y-chromosome sequences reveal an extremely recent origin of the most common North African paternal lineage E-M183 (M81). Scientific reports 7, 15941 (2017). 10.1038/s41598-017-16271-y

36 Semino, O. et al. Origin, Diffusion, and Differentiation of Y-Chromosome Haplogroups E and J: Inferences on the Neolithization of Europe and Later Migratory Events in the Mediterranean Area. The American Journal of Human Genetics 74, 1023–1034 (2004). 10.1086/386295

37 Vezzosi, A., and A. Sabato. The New Genealogical Tree of the Da Vinci Family for Leonardo’s DNA. Ancestors and Descendants in Direct Male Line down to the Present XXI Generation. Human Evolution 36 (2021).

38 da Vinci, L. The Heart, Bronchi, and Bronchial Vessels. Academic Medicine 96, 516 (2021). 10.1097/acm.0000000000003380

39 Sweet, D., Lorente, M., Lorente, J. A., Valenzuela, A. & Villanueva, E. An improved method to recover saliva from human skin: the double swab technique. J Forensic Sci 42, 320–322 (1997).

40 Torralba, M. G., Kuelbs, C., Moncera, K. J., Roby, R. & Nelson, K. E. Characterizing Microbial Signatures on Sculptures and Paintings of Similar Provenance. Microb Ecol 81, 1098–1105 (2021). 10.1007/s00248-020-01504-x

41 Nurk, S., Meleshko, D., Korobeynikov, A. & Pevzner, P. A. metaSPAdes: a new versatile metagenomic assembler. Genome research 27, 824–834 (2017).

42 Shen, W., Le, S., Li, Y. & Hu, F. SeqKit: a cross-platform and ultrafast toolkit for FASTA/Q file manipulation. PloS one 11, e0163962 (2016).

43 Langmead, B. & Salzberg, S. L. Fast gapped-read alignment with Bowtie 2. Nature methods 9, 357–359 (2012).

44 Li, H. et al. The sequence alignment/map format and SAMtools. bioinformatics 25, 2078–2079 (2009).

45 Liao, Y., Smyth, G. K. & Shi, W. featureCounts: an efficient general purpose program for assigning sequence reads to genomic features. Bioinformatics 30, 923–930 (2014). 10.1093/bioinformatics/btt656

46 Hyatt, D. et al. Prodigal: prokaryotic gene recognition and translation initiation site identification. BMC Bioinformatics 11, 119 (2010). 10.1186/1471-2105-11-119

47 von Meijenfeldt, F. A. B., Arkhipova, K., Cambuy, D. D., Coutinho, F. H. & Dutilh, B. E. Robust taxonomic classification of uncharted microbial sequences and bins with CAT and BAT. Genome Biol 20, 217 (2019). 10.1186/s13059-019-1817-x

48 Li, H. & Durbin, R. Fast and accurate short read alignment with Burrows-Wheeler transform. Bioinformatics 25, 1754–1760 (2009). 10.1093/bioinformatics/btp324

49 Danecek, P. et al. Twelve years of SAMtools and BCFtools. Gigascience 10 (2021). 10.1093/gigascience/giab008

50 Poznik, G. D. et al. Sequencing Y chromosomes resolves discrepancy in time to common ancestor of males versus females. Science 341, 562–565 (2013). 10.1126/science.1237619

51 Waskom, M. L. Seaborn: statistical data visualization. Journal of open source software 6, 3021 (2021).

