## Supplemental Figures for "Biological signatures of history: Examination of composite biomes and Y chromosome analysis from da Vinci-associated cultural artifacts"

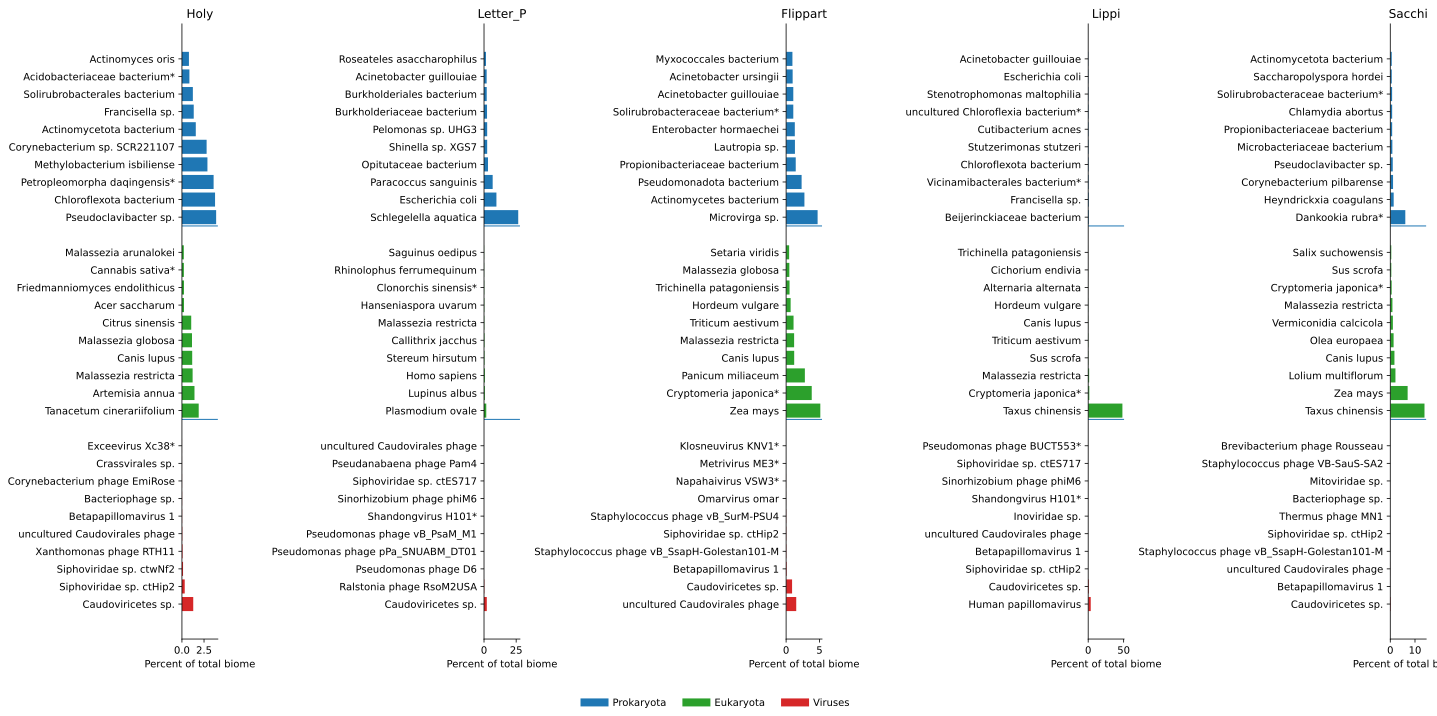

**Fig. S1: Top ten species for prokaryote, eukaryote and viruses.**

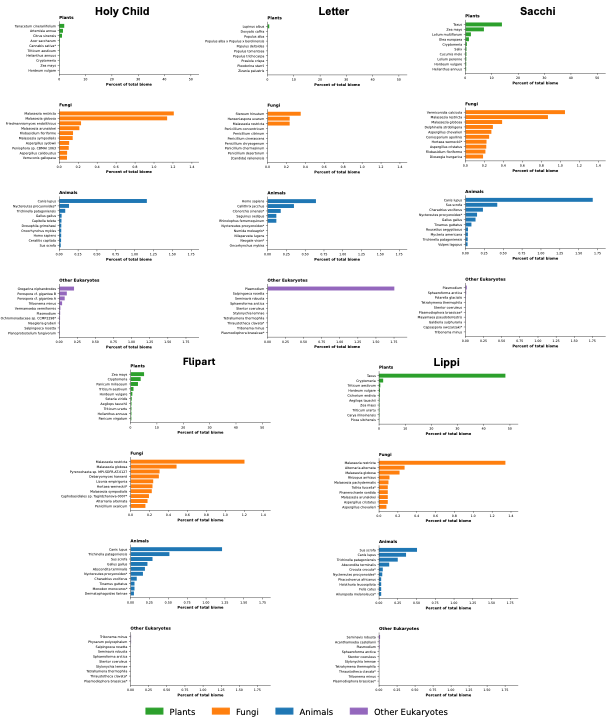

**Fig S2. Top ten species of plants, fungi, animal and other eukaryotes in samples.**

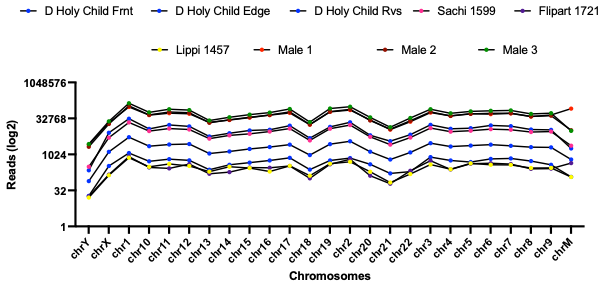

**Fig S3. Human DNA mapping to specific chromosomes.**

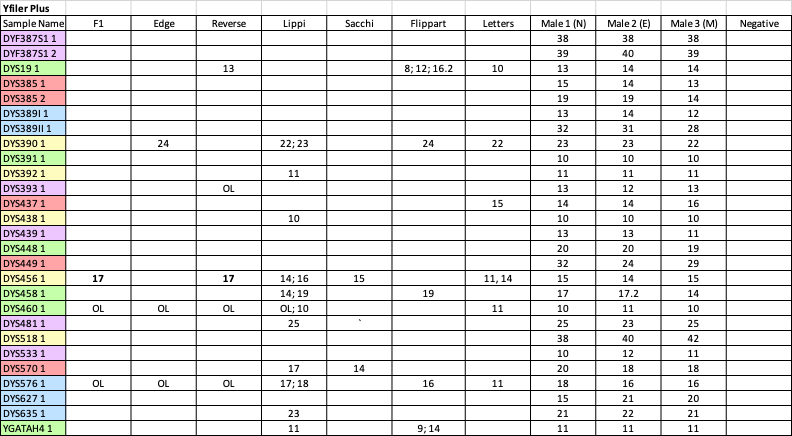

**Fig. S4 Y filer results.**
